# scATAnno: Automated Cell Type Annotation for single-cell ATAC Sequencing Data

**DOI:** 10.1101/2023.06.01.543296

**Authors:** Yijia Jiang, Zhirui Hu, Allen W. Lynch, Junchen Jiang, Alexander Zhu, Ziqi Zeng, Yi Zhang, Gongwei Wu, Yingtian Xie, Rong Li, Ningxuan Zhou, Cliff Meyer, Paloma Cejas, Myles Brown, Henry W. Long, Xintao Qiu

## Abstract

Recent advances in single-cell epigenomic techniques have created a growing demand for scATAC-seq analysis. One key analysis task is to determine cell type identity based on the epigenetic data. We introduce scATAnno, a python package designed to automatically annotate scATAC-seq data using large-scale scATAC-seq reference atlases. This workflow generates the reference atlases from publicly available datasets enabling accurate cell type annotation by integrating query data with reference atlases, without the use of scRNA-seq data. To enhance annotation accuracy, we have incorporated KNN-based and weighted distance-based uncertainty scores to effectively detect cell populations within the query data that are distinct from all cell types in the reference data. We compare and benchmark scATAnno against 7 other published approaches for cell annotation and show superior performance in multiple data sets and metrics. We showcase the utility of scATAnno across multiple datasets, including peripheral blood mononuclear cell (PBMC), Triple Negative Breast Cancer (TNBC), and basal cell carcinoma (BCC), and demonstrate that scATAnno accurately annotates cell types across conditions. Overall, scATAnno is a useful tool for scATAC-seq reference building and cell type annotation in scATAC-seq data and can aid in the interpretation of new scATAC-seq datasets in complex biological systems.

## Introduction

Recent advances in single-cell technologies enable the profiling of cellular heterogeneity in complex tissues at single-cell resolution. The single cell assay for transposase-accessible chromatin using sequencing (scATAC-seq) profiles chromatin accessibility in single cells and uncovers regulatory elements, known as cis-regulatory elements (CREs), which govern cell-type-specific gene expression patterns [1, 2]. Thus, the scATAC-seq profile plays an important role in understanding transcriptional patterns and chromatin configurations underling cell states.

To identify what cell types are present in single-cell data, cell type annotation methods are needed. Many existing methods were developed for single-cell RNA-seq (scRNA-seq) data. These methods can be broadly categorized into two types: 1) leveraging well-annotated reference atlases to label cells in new datasets (known as query data) and 2) using a set of known gene markers to infer cell types. The first approach is used by tools such as SingleR, which predicts unannotated cells based on gene expression correlation with reference [3], and scANVI, which is a deep generative model to transfer labels from reference to query data [4], while the second is exemplified by Garnett, which annotates cells based on pre-defined cell type-specific markers [5]. Although many methods are designed for scRNA-seq data, there are a limited number of suitable tools to automatically annotate scATAC-seq data.

Single-cell chromatin accessibility data present unique challenges compared to the scRNA-seq data. The first challenge is data sparsity and the high dimensionality of epigenomic features. In contrast to scRNA-seq data, which typically contain one to several hundred transcripts per gene, scATAC-seq data often has a few fragment counts per peak, due to the presence of two copies of DNA in most cells at most locations, resulting in data sparsity [1, 2]. Moreover, scATAC-seq has much higher feature dimensions, as the number of potential genomic regulatory regions could go up to millions, while the human genome only contains approximately 20,000 coding genes [6]. Another challenge is the lack of marker enhancers to annotate scATAC-seq data. Although previous efforts profiled regulatory elements in tissues and cell lines [7], there is still a lack of complete catalogs of cell-type specific CREs [2]. Despite these challenges, it is important to annotate cell types at the scATAC level because the epigenetic landscape governs gene expression and plays an important role in determining cell types. Most of the existing scATAC-seq annotation methods rely on converting peaks or genomic regions to gene activity scores and then employ various approaches to determine cell type. These approaches include 1) using scRNA-seq-specific methods to predict cell types, such as Seurat v3 [8], 2) utilizing supervised learning models to train on scATAC-seq reference and infer cell types of query data, such as Cellcano [9], or 3) annotating cell type labels based on gene markers, such as MAESTRO [10]. However, such methods rely on promoter and gene body regions as proxies for computing gene activity scores, and do not fully take advantage of hundreds of thousands open regions at enhancers that are distant from promoters in the scATAC-seq profile [11]. There is a demand to better use CREs in the enhancer regions for cell type annotation at single-cell level. The development of single-cell epigenomic techniques has also led to increased availability of large-scale scATAC-seq datasets and a larger number of cell-type-specific regulatory elements, spanning from healthy adult cells to tumor-infiltrating lymphocytes [12–14]. These scATAC-seq reference atlases collect data from different tissues and conditions and enable the construction of peak references by aggregating robust signals from a sufficient number of single cells providing an opportunity to automate cell type annotation for scATAC-seq data.

We present scATAnno, a Python-based workflow that directly and automatically annotates scATAC-seq data based on scATAC-seq reference atlases. Unlike many existing approaches, scATAnno directly uses peaks or CRE genomic regions as input features, eliminating the need to convert the epigenomic features into estimated gene activity scores. scATAnno tackles the high dimensionality of scATAC-seq data by leveraging spectral embedding to efficiently transform the data into a low dimensional space. To create catalogs of CREs at single-cell level, scATAnno uses chromatin state profiles from a large-scale reference atlas to generate peak signals and reference peaks.

We show that the scATAnno workflow can generate scATAC-seq reference atlases from publicly available datasets and various scATAC-seq technologies, including 10x Genomics [13] and combinatorial indexing approaches [12, 15]. Additionally, users can readily build customized scATAC-seq reference atlas using scATAnno workflow (Methods). scATAnno enables accurate cell type annotation by integrating query data with reference atlases in an unsupervised manner. The results are further augmented with two uncertainty score metrics to quantitatively assess the confidence of cell-type assignment. The first uncertainty score is based on the K-Nearest Neighbor (KNN) and determines if there is just one cell type most similar to the query cell. The second uncertainty score is derived from a novel computation of the weighted distance between the query cell and reference cell type centroids in a low dimensional space. This score determines if the one most similar cell type is “close enough” to the query cell to be assigned confidently. We compare and benchmark scATAnno against 7 other published approaches for cell annotation and show superior performance in multiple data sets and metrics. We showcase the utility of scATAnno in multiple case studies, including peripheral blood mononuclear cell (PBMC), Triple Negative Breast Cancer (TNBC), and basal cell carcinoma (BCC), and demonstrate that scATAnno accurately annotates cell types and identifies unknown cell populations not represented in the reference atlas. Furthermore, scATAnno enables efficient cell type annotation across tumor conditions and accurately annotates tumor-infiltrating lymphocytes (TIL) in TNBC using a BCC TIL atlas. Overall, scATAnno is a useful cell-type annotation tool specifically designed for scATAC-seq data and expedites our understanding of chromatin-based cellular heterogeneity in complex biological systems.

## Results

### The scATAnno workflow

scATAnno is a powerful automated tool to annotate cell types for scATACseq data. It takes fragment files of the query data and a reference atlas of choice as input and predicts query cell types at both the individual cell and cell cluster levels. As shown in Figure 1, scATAnno first constructs peak-by-cell matrices of the reference atlas and the query data (Methods). Reference atlases are generated from publicly available datasets, and the feature space is obtained either from the original publication [12] or through MACS2 peak-calling algorithm [13, 16]. If users want to build their own reference atlas data, it is easy to build customized reference atlases following the reference building steps (Methods). Unlike scRNA-seq data, scATAC-seq lacks marker peaks, and thus the feature space varies between scATAC-seq datasets. To unify the feature space, scATAnno uses the set of peaks from the reference atlas and counts reads from the query dataset that fall within these peaks, generating a peak-by-cell matrix for the query dataset (methods). In step two, matrices of reference and query cells are concatenated, followed by dimensionality reduction using spectral embedding (Methods). The workflow of scATAnno incorporates the SnapATAC2 tool into the workflow to perform dimensionality reduction [17, 18]. This converts the concatenated count matrix into a cell-cell similarity matrix and then performs eigenvector decomposition (Methods). From a recent paper, SnapATAC2 shows superior performance in dimensionality reduction, achieving shorter runtime, memory usage and better clustering results than existing dimensionality reduction methods. We therefore decided to use this approach to reduce the dimensionality for the integrated reference and query data in our workflow [17, 19]. In step three, following the projection of reference and query cells into the same low dimensional space, scATAnno optionally employs a batch removal method [20], to mitigate any potential batch effects, such as technical variations, between the reference and query data.

**Figure 1:**
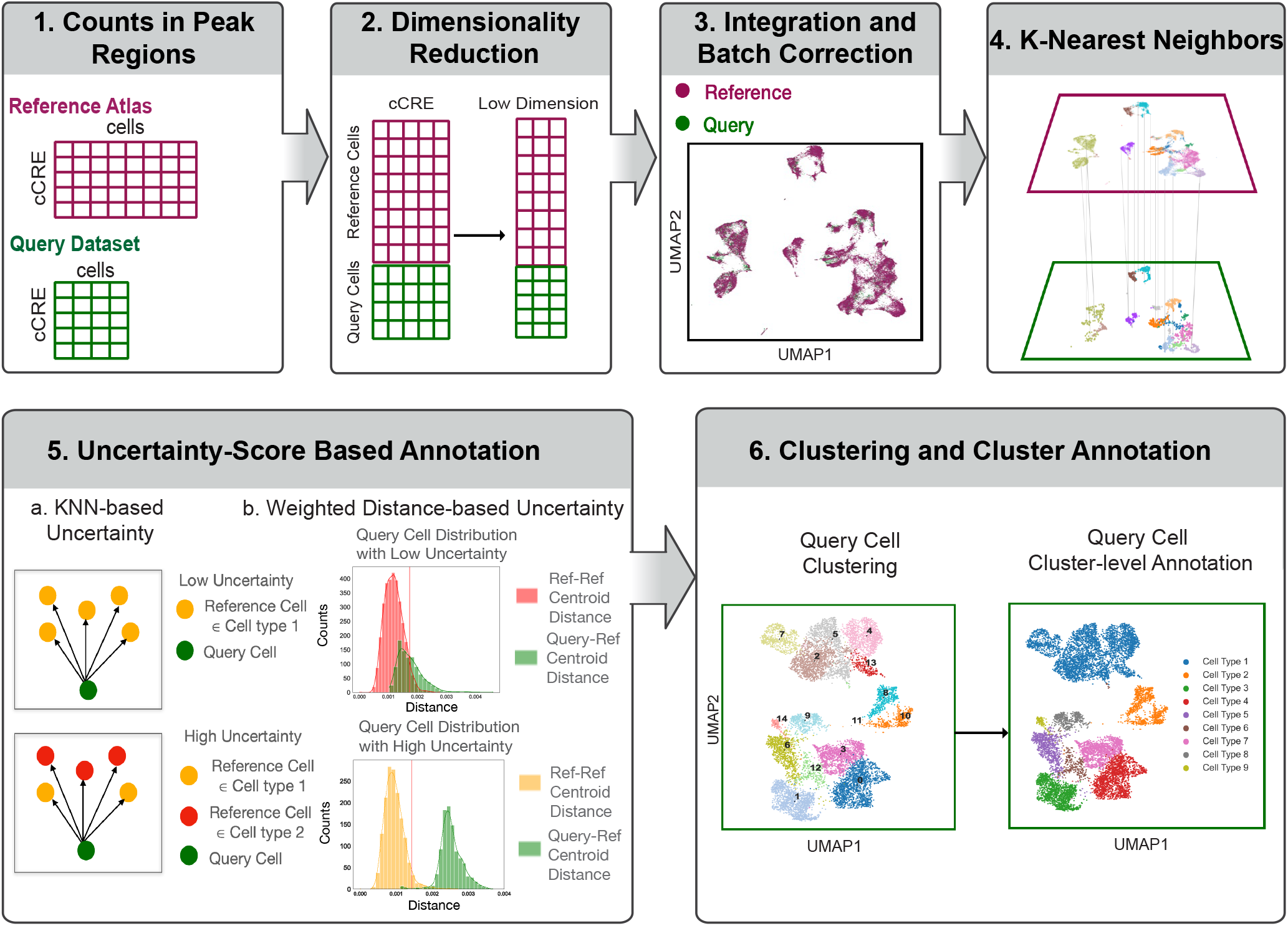
Schematic Overview of scATAnno. scATAnno consists of six steps: 1) peak-by-cell matrix construction of reference atlas and query data, 2) dimensionality reduction using spectral embedding, 3) integration and batch effect removal, 4) KNN cell-type assignment. The lower green plane shows query cells, each annotated using its closest neighbors in the reference cells in the upper purple plane, 5) Two uncertainty score measurements: KNNbased (panel a) and weighted distance-based (panel b). In the KNN-based uncertainty (panel a), a query cell is either assigned low uncertainty score if its K-nearest neighbors belong to the same reference cell type (top) or assigned high uncertainty score if its K-nearest neighbors belong to mixed reference cell types (bottom). In the weighted distance-based uncertainty (panel b), the distances of query cells to the centroid of each reference cell type (Query-Ref Centroid Distance) are compared with the within-cell type distribution of each reference cell type (Ref-Ref Centroid Distance). Query cells that fall within 95 percentile of the within-cell type distribution are assigned low uncertainty scores (top), and query cells that fall outside of 95 percentile of the within-cell type distribution are assigned high uncertainty scores (bottom). The red vertical line in panel b represent 95 percentile of Ref-Ref Centroid Distance distribution, 6) clustering and annotation of query cells at the cell cluster level

In the fourth step, query cells are assigned cell labels using K-nearest neighbors (KNN) approach. Each query cell is annotated based on its K closest neighbors in the reference atlas. To measure the confidence of the KNN assignment, two distinct uncertainty score metrics are utilized in the fifth step. The first metric is the KNN-based uncertainty score, which was described in previous publications [21]. This score is computed by considering the closest neighbors of a query cell in the reference atlas. Query cells with nearest neighbors predominantly belonging to a single cell type are assigned low uncertainty scores, while those with neighbors consisting of a mixture of cell types receive higher uncertainty scores (Methods). While useful, the KNN-based uncertainty score has limitations and does not reflect the distance between a query cell and its assigned cell type cluster. This becomes a problem when query data contains an unknown cell population not represented in the reference, leading to inaccurate cell type assignment. To address this limitation, we introduce a novel computation of a weighted distance-based uncertainty score, based on the intuition that query cells located far from the centroid of an assigned cell type represent an unknown population. The computation takes three steps. First, we identify centroids of reference cell types based on low dimensional components. Second, we hypothesize that low dimensional components have varying discriminate power for a specific cell type, so for each cell type, we assign different weights to the components based on the closeness of reference cells to their respective cell type’s centroid along each dimension (methods). Components with higher weights indicate greater importance in distinguishing a cell type. Third, a reference distance distribution is generated for each cell type by calculating the weighted distance between each reference cell and the centroid of its cell type. Subsequently, we compute the weighted distance of a query cell to the centroid of its assigned cell type and compare the query distance with the reference distance distribution. If the query-ref centroid distance is greater than 95th percentile of the reference distance distribution, a query cell is assigned a high uncertainty score. The final uncertainty score for a query cell is determined by taking the maximum value between the KNN-based and weighted distance-based uncertainty scores. Query cells with high uncertainty scores are considered as unknown cells. Finally, in step six, query cells are clustered using the Leiden algorithm [22] and each cell cluster is annotated based on the most abundant cell type present in the cluster.

### scATAnno builds a reference atlas for healthy adult cells and achieves accurate cell type annotation

We first test the performance of scATAnno by annotating cells within the reference dataset. We built a healthy adult reference atlas across 30 tissues from a publicly available dataset [12]. Specifically, we selected 100,158 deep-sequenced cells, and 890,130 adult-specific peaks provided by the original study (methods). scATAnno builds a ready-to-use reference atlas with 28 well-separated cell types (Supplementary Figure 1A). To assess the accuracy of scATAnno in predicting cell type, we utilized cells generated from the original study but excluded from our reference building as query cells. scATAnno successfully integrates the query cells with the healthy adult reference atlas, as shown by UMAP (Supplementary Figure 1B). We further visualized predicted cell types of query cells along with their associated uncertainty scores in Supplementary Figure 1D. Comparing the annotations from the original paper (Supplementary Figure 1C), scATAnno demonstrates accurate classification of most cells and displays high uncertainty scores for individual cells situated in transition regions between cell clusters and peripheral regions of any cluster (Supplementary Figure 1D). At the cluster-level annotation, scATAnno achieved an accuracy of 98.4% and an average F1 score of 98.2% (Supplementary Figure 1F) (method). We also examined the impact of cell type similarities on performance by analyzing the accuracy and weighted F1 scores for each cell type. Notably, cell types that exhibit similarity to other cell types, such as the Ionic mesenchymal cells, displayed relatively lower accuracy and F1 scores (Supplementary Figure 1E). Additionally, we compared the annotation performance using all peak regions versus promoter regions and found that by including all peak regions, the annotation performance is greatly improved compared to using promoter regions alone. In Supplementary Figure 1F, we intersected all peak regions with TSS promoter regions extending to 1kb, 2kb, 5kb and 10kb, and compared the annotation accuracy using extended promoter regions versus all regions. We observed that as the number of promoter regions increased, the accuracy also increased. Moreover, using all regions significantly enhanced the cell type annotation accuracy (98%) compared to the highest accuracy using promoter regions extended 10kb (77%) (Supplementary Figure 1F). Further breaking down the accuracy score of each cell type, we consistently found that using all regions achieved higher accuracy score for each cell type than using promoter regions only (Supplementary Figure 1G). Several cell types, such as Ionic Mesenchymal, Keratinocyte, Luteal, and PNC-derived cells, consistently exhibited poor performance when using promoter regions only. This suggests that relying solely on promoter regions may fail to distinguish certain celltypes.

Next, we investigated how accurately scATAnno detects an unknown cell population in query data not represented in the reference atlas. We explore the annotation in two scenarios: 1) ablate a cell type that is similar to other cell types, and 2) ablate a cell type distinct from other cell types (Supplementary Figure 2A-C). We first ablated the vascular smooth muscle cell type from the reference atlas to mimic the unknown cell-type scenario 1 (Supplementary Figure 2B). The original reference atlas (Supplementary Figure 2A) demonstrates the close clustering of the vascular smooth muscle cell type with the mural cell type, indicating cell type similarity. Next, we show step-by-step annotation results for query data visualized by UMAP (Supplementary Figure 2D), highlighting the vascular smooth muscle cells in query data in the dashed circle. We expect a high proportion of vascular smooth muscle cells in query data is unknown since this cell type is removed from the reference atlas. In the KNN assignment step, more than 80% vascular smooth muscle cells in query data are assigned high uncertainty scores and successfully detected as unknown. This outcome could be attributed to the mixed KNN neighbors of these query cells. As most vascular smooth muscle query cells are detected by the KNN-based uncertainty score, the weighted distance-based uncertainty score does not significantly improve the result (Supplementary Figure 2E). At the cluster level, all vascular smooth muscle cells in query data are correctly annotated as unknown cells.

In the second scenario, we ablated the Adrenal Cortical cell type from the reference atlas (Supplementary Figure 2C) and presented prediction results for the query data visualized by UMAP, highlighting the adrenal cortical cells in query data in the dashed circle (Supplementary Figure 2F). The Adrenal Cortical cell type is distinct from other cell types as it is well-separated from any other cell type (Supplementary Figure 2A). In the KNN assignment step, most adrenal cortical cells in query data are incorrectly annotated as immune or mesenchymal cells with low uncertainty scores (Supplementary Figure 2F). The misclassification is due to the fact that when the ablated cell type is distinct from other cell types, the KNN assignment erroneously assigns query cells to the next closest reference cluster while displaying low uncertainty scores. To overcome this limitation, we introduced the weighted distance-based uncertainty score. After applying this weighted distance-based uncertainty, we observe a significant increase in the proportion of expected unknown cells from 7.3% to 66.0%, and all adrenal cortical cells are correctly identified as unknown at the cluster level (Supplementary Figure 2G). This demonstrates that the utilization of this more sophisticated weighted distancebased score results in a substantial improvement in the detection of unknown cell populations in the query data.

### scATAnno builds a reference atlas for PBMC immune cells and enables accurate classification of immune cell subtypes in PBMC Multiome data

We showcase the utility of scATAnno in integrating data from multiple datasets and cell-type annotation by applying it to PBMC data. We built a large-scale PBMC reference atlas from publicly available data [13]. To construct this atlas, we downloaded raw FASTQ files to generate BAM files and fragment files across 15 healthy individuals and sampled cells. We then selected the immune cells (39441 cells) and generated a reference peak set (344492 peaks) by using MACS2 to call peaks (methods). The UMAP plot displays a ready-to-use PBMC reference atlas encompassing 14 cell types in Supplementary Figure 3A. To demonstrate the inter-dataset annotation performance of scATAnno, we downloaded PBMC 10K Multiome data from 10x Genomics as query cells (methods). The advantage of using the Multiome data is the availability of both scATAC-seq and scRNA-seq profiles, which help validate the cell-type annotation results using peak signals and gene expression patterns of canonical markers from the same cells. As shown in Supplementary Figure 3B, scATAnno successfully integrates and removes the batch effects of the PBMC reference atlas and the query data.

To evaluate the performance of scATAnno, we compare the predicted cell types with Seurat v3 [8] which first converts scATAC-seq data to gene activity scores and transfers labels from the scRNA-seq profile to scATAC-seq data. The Seurat v3 annotation is directly obtained from the built-in data object stored in the Seurat function (methods). Most cell types predicted by scATAnno are in agreement with that by Seurat v3, except for the two B cell subtypes (Figure 2A-C). scATAnno annotates the lower B cell cluster as Naive B and the upper cluster as Memory B, while Seurat v3 annotates the opposite. To validate annotations produced by scATAnno, we leveraged the scRNA-seq profile and selected gene markers to distinguish two B cell subtypes. Specifically, CD27 and TNFRSF13B genes are selected as Memory B markers, and IGHM and TCL1A are chosen as Naive B markers [23]. In Figure 2D, Naive B cell markers (CD27 and TNFRSF13B) exhibit increased expression levels in the cell cluster annotated as Naive B, and Memory B cell markers are upregulated in the cell cluster annotated as Memory B, confirming the cell identities predicted by scATAnno. Lastly, scATAnno detects unknown cells not represented in the PBMC reference atlas, for example, a group of HSPC cells annotated by Seurat v3 is classified as unknown cells, due to the lack of this cell type in the PBMC reference atlas.

**Figure 2:**
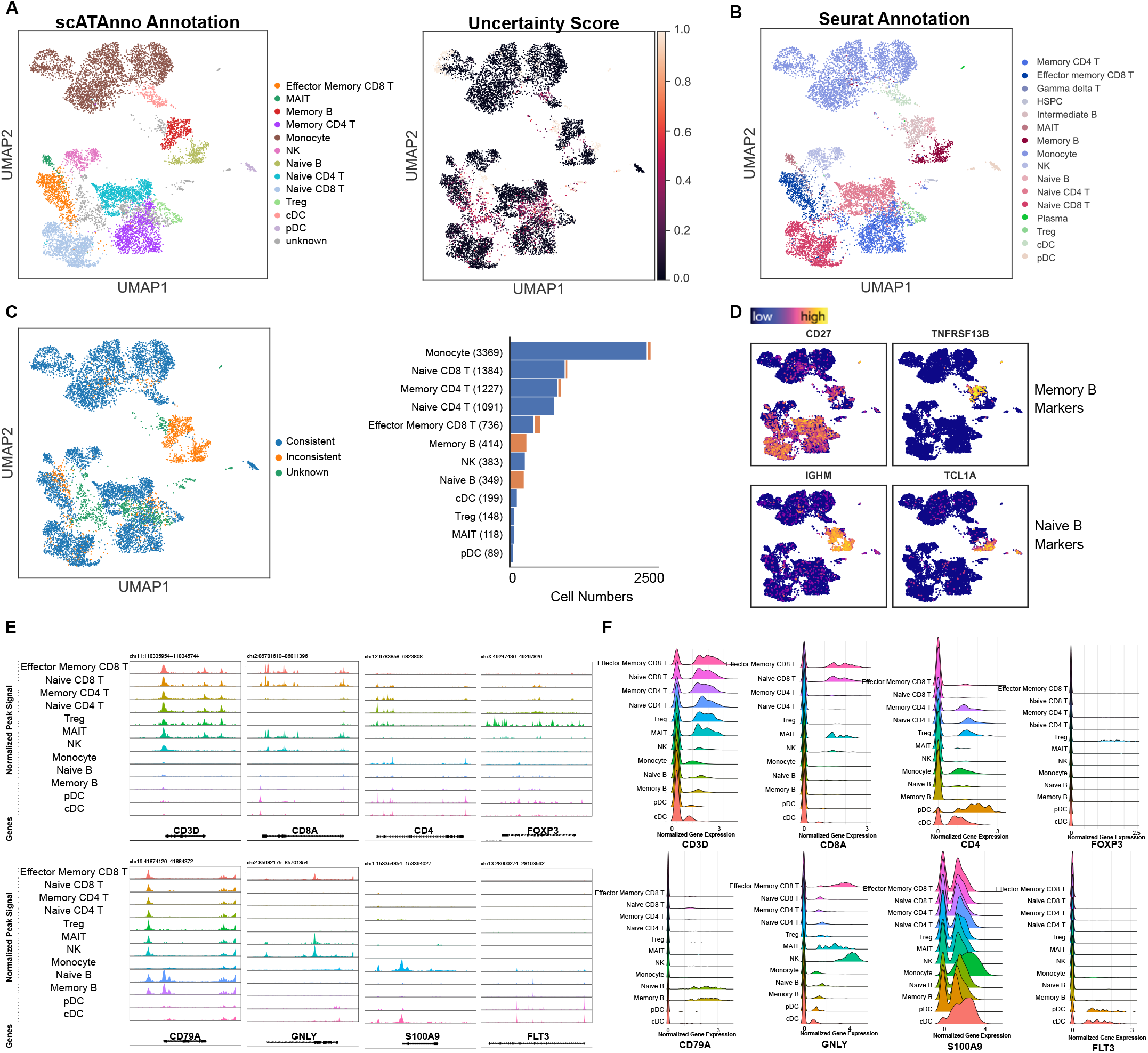
PBMC Case Study. (A) UMAP representation of query cells colored by cell types predicted from scATAnno (left) and distribution of uncertainty score (right) (B) UMAP representation of query cells colored by cell types predicted from Seurat annotation (C) Comparisons of cell type annotation from scATAnno and Seurat shown by UMAP (left), and by stacked bar plot of predicted cell types (right). The stacked bar plot displays the counts of consistently annotated cells, inconsistently annotated cells, and unknown cells for each cell type (D) Gene expression levels of signature markers for B cell subtypes using scRNA-profile, (E) Coverage plots of chromatin accessibility over signature genes across predicted cell types: CD3D for T cells, CD8A for CD8 T, CD4 for CD4 T, FOXP3 for Treg, CD79A for B cells, GNLY for NK, S100A9 for Monocytes, and FLT3 for cDC and pDC (F) Ridge plots of the gene expression levels of signature genes

To further validate cell identities yielded by scATAnno, we examined the chromatin accessibility and gene expression of signature genes by plotting the peak signals using the scATAC-seq profile and gene expression distribution using the scRNA-seq profile. The coverage plot (Figure 2E) shows the normalized peak signal over a given genomic region ±3kb, while the ridge plot (Figure 2F) shows the distribution of normalized gene expression across cell groups. As shown in Figure 2E, the epigenetic features in the scATAC-seq profile display specific enrichment of peaks in corresponding cell types. For instance, T cell subtypes, including CD8 T subtypes, CD4 T subtypes, and Treg, exhibit enrichment of peaks over the CD3D gene, which is a canonical marker of T cells. Specifically, CD8 and CD4 T cell subtypes show enriched peak signals over CD8A and CD4 genomic regions, respectively. Regulatory T cells (Tregs) show specific peak enrichment over the FOXP3 locus, a gene marker for regulatory T cells. The gene expression patterns of signature genes in the scRNA-seq profile further confirm the cell type annotation result (Figure 2F). For example, the CD3D gene, a marker for T cells, exhibits a high expression level in T cell subtypes. CD8A and CD4 genes show increased gene expression levels in the respective CD8 and CD4 T cells. FOXP3 gene shows an increased expression level in the Treg population, although we observe more distinct gene activity in the scATAC profile than scRNA profile. Finally, CD79A, GNLY, S100A9, and FLT3 show strong peak signals and gene expression levels in B cells, NK cells, monocytes, and dendritic cells, respectively. Additionally, we examined chromatin accessibility over TRAV1-2, SLCA410, and PRSS35 genes, which are known as markers for MAIT cells [23, 24] (Supplementary Figure 3C). The enhanced chromatin accessibility over these regions confirms MAIT cell identity. The epigenetic features and gene expression patterns of signature genes support cell-type annotation produced by scATAnno, demonstrating its robust capability in correctly identifying B cell subtypes misannotated by Seurat v3.

### Benchmarking scATAnno against existing scATAC-seq cell typing methods

scATAnno stands out with distinctive features for annotating scATAC-seq cell types compared to other existing methods, as outlined in Table 1. While only scATAnno, Cellcano [25] and Signac [24] are designed for scATAC-seq anotation, Seurat [8], scJoint [26], singleR [3] and scID [27] are developed for scRNA-seq data. Notably, scATAnno is unique in its capability to harness enhancer signals, eliminating the need to translate peaks into proximal gene activity scores. For procedures that demand a gene activity score input, we adhered to each method’s guidelines to derive the gene activity score matrix. For tools intended for scRNA-seq data, the gene activity score was first procured from ArchR, followed by annotation based on a reference atlas and gene markers. Regarding uncertainty-aware mechanisms, scATAnno, Seurat and Signac can yield uncertainty scores to predict unknown cells, while the other methods do not provide a means to quantify the reliability of their annotation results (Table 1).

To benchmark scATAnno against Cellcano, scJoint, Signac, Seurat, singleR and scID, we used two distinct datasets to evaluate performance. The first dataset is derived from a lymph node tumor of a patient diagnosed with B cell Lymphoma, and the data can be downloaded from 10X (Method). The second dataset features PBMC cells isolated through Fluorescence-activated cell sorting (FACS) (GSE123578).

We selected these datasets as they are accompanied by high-confidence cell type labels. We first benchmarked all methods on scATAC-seq data from a single-cell multi-omic dataset of a B cell lymphoma patient to investigate the performance of annotation in a more real-world disease context. This dataset has well-defined ground truth cell labels based on the matched scRNA-seq data from the same nuclei and is composed of immune cells and tumor B cells. Though similar cell type signature genes are enriched in malignant cells to normal cells, the tumor cells are often harboring cytogenetic aberrations, leading to distinguished cell status and types. Therefore, tumor B cells are distinct from normal B cells. We used the healthy PBMC reference atlas for all methods, using this B Lymphoma dataset as query. Supplementary Figure 4A shows UMAP of cells colored by cell types annotated by scATAnno, Cellcano, and Signac. Comparing annotations with ground truth labels, scATAnno successfully identifies the majority of tumor B cells as unknown cells, whereas Cellcano annotates B cells as a mixed cluster of Monocytes and B cells, and Signac fails to distinguish tumor B cells from normal B cells. Comparing accuracy scores across seven methods shows that scATAnno achieves the highest accuracy 0.90 followed by Signac with accuracy 0.715, and Cellcano with accuracy 0.581 (Supplementary Figure 4B). scATAnno significantly outperforms the other methods in this case study. In this regard, unlike other conventional methods counting B-cell lymphoma cells as B cells, scATAnno correctly identified the lymphoma cells as a distinct cell population, which is consistent with scRNA-seq analysis (Supplementary Figure 4D).

Our second benchmark used FACS PBMC data as query data since it has very high quality cell types. We downloaded raw fragment files, used Signac to filter out low quality cells (Method), and ended up with 15,643 immune cells, composed of CD8 T cells, CD4 T cells, B cells, Monocytes and NK cells. We applied all annotation methods on this dataset and compared with ground truth labels determined by the cell sorting. We created five annotation tasks by using all reference cells, or systemtically removing each cell type, CD8 T cells, CD4 T cells, B cells, Monocytes and NK cell types, from the PBMC reference atlas, to test whether ablations of cell types in the reference atlas would affect the performance of annotation (Methods). Compared the average accuracy score across methods, scATAnno achieves the best average accuracy score with 0.811; Cellcano is the second with average accuracy as 0.769; and scJoint is the third with average accuracy 0.742 (Supplementary Figure 4E-G). scATAnno significantly outperformed the other methods in the ablation study, and it is able to identify the unknown cell population ablated from the reference atlas (Supplementary Figure 4F). We conclude scATAnno performs the best and is the most robust method for cell type annotation in PBMC sorted data.

### scATAnno identifies cancer cell cluster and normal cells in the microenvironement

Cancer samples typically consist of tumor cells and immune cells. To identify tumor cells and immune cells in the tumor microenvironment (TME), we next show the utility of combinations of reference atlases including both healthy adult cells and cancer cells atlas to enable multi-layer cell-type annotation in the two following sections.

First, we use scATAnno to distinguish tumor cells from immune cells in cancer samples using a healthy adult reference atlas. We selected two BCC samples held out from the BCC TIL reference building as query data and mapped them to the healthy adult reference atlas. The BCC samples are preor post-treated by immunotherapy and comprise a diverse ecosystem of cell types in the TME. To balance the treatment condition, we deliberately selected one pre-treatment and one post-treatment sample as the query. The query data contains tumor cells that are not present in the healthy adult reference atlas; therefore, intuitively the tumor cells should display high uncertainty scores and be identified as unknown. As shown in Supplementary Figure 5A-B, query cells are successfully integrated with the reference atlas. By scrutinizing the predicted cell types and uncertainty score distribution on the UMAP space of query cells (Figure 3A-B), we observed that tumor cells show high uncertainty scores and are accurately annotated as unknown at the cluster level. Furthermore, scATAnno accurately annotates endothelial and immune cells with low uncertainty scores (Figure 3B). Fibroblasts on the other hand are identified as mural cells, also known as pericytes [12, 28], which could be a result of the strong connection between pericytes and fibroblasts in TME [28, 29]. The chromatin accessibility profile confirms the enrichment of peak signal over PDGFRB, an important marker of pericytes [30], in the mural group (Supplementary Figure 5D). scATAnno can leverage the healthy adult reference atlas to annotate cell types in tumor samples and accurately identify the normal cells, immune cells while annotating tumor cells as unknown.

**Figure 3:**
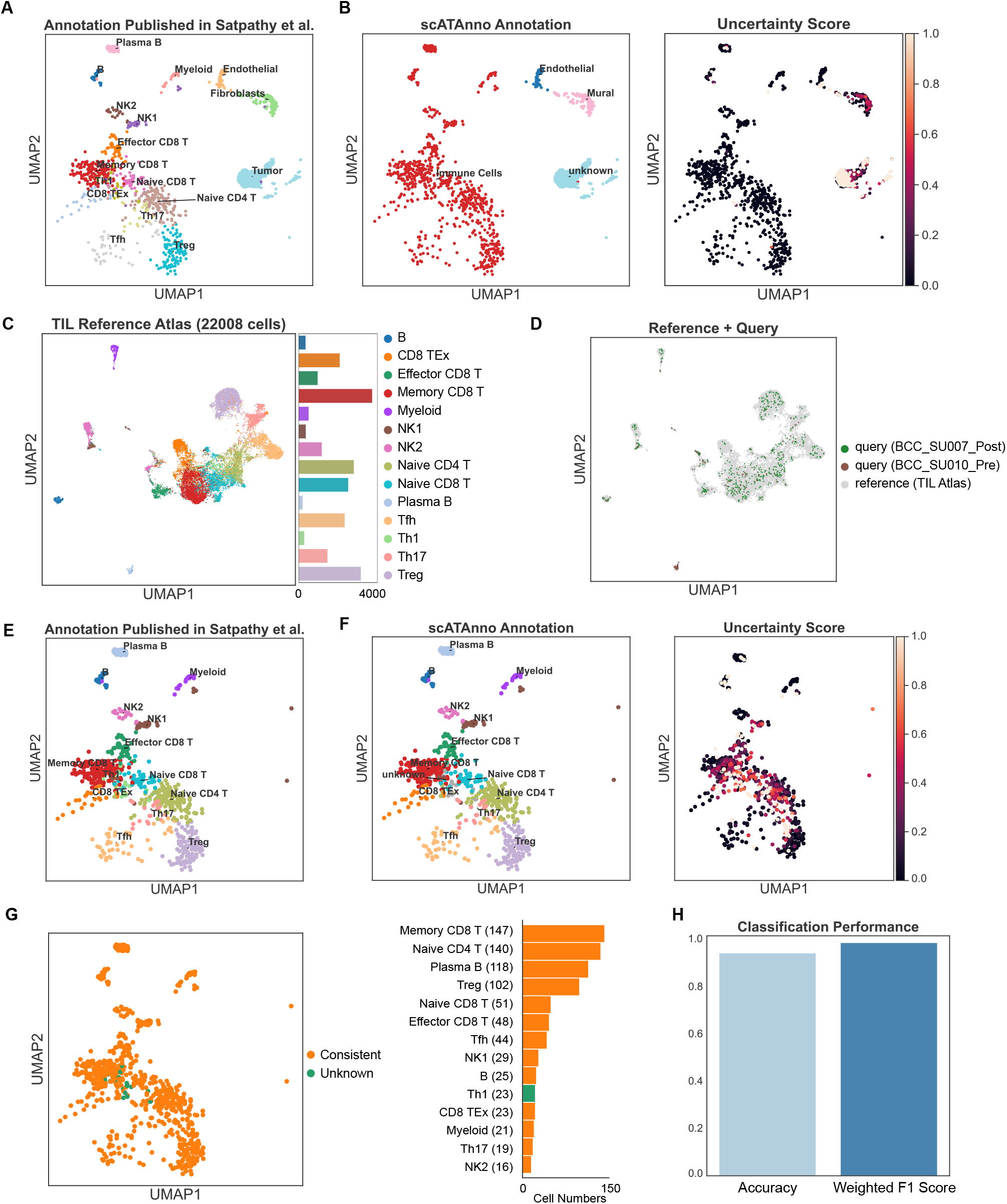
BCC Case Study. (A) UMAP representation of query cells colored by cell types from the original publication (Sapathy et al.) (B) UMAP representation of query cells colored by cell types predicted from scATAnno (left) and distribution of uncertainty score (right) (C) Construction of BCC TIL reference atlas (D) Integration of immune cells from two holdout BCC samples and TIL reference atlas (E) UMAP representation of immune cells from the query data colored by cell types from the original publication (Sapathy et al.) (F) UMAP representation of immune cells colored by cell types predicted from scATAnno along with the uncertainty score distribution (G) Comparison of cell type annotations between scATAnno and original study shown by UMAP (left), by stacked bar plot (right) of cell numbers. The stacked bar plot displays the counts of consistently annotated cells and unknown cells for each cell type H) Classification accuracy and weighted F1 score

**Figure 4:**
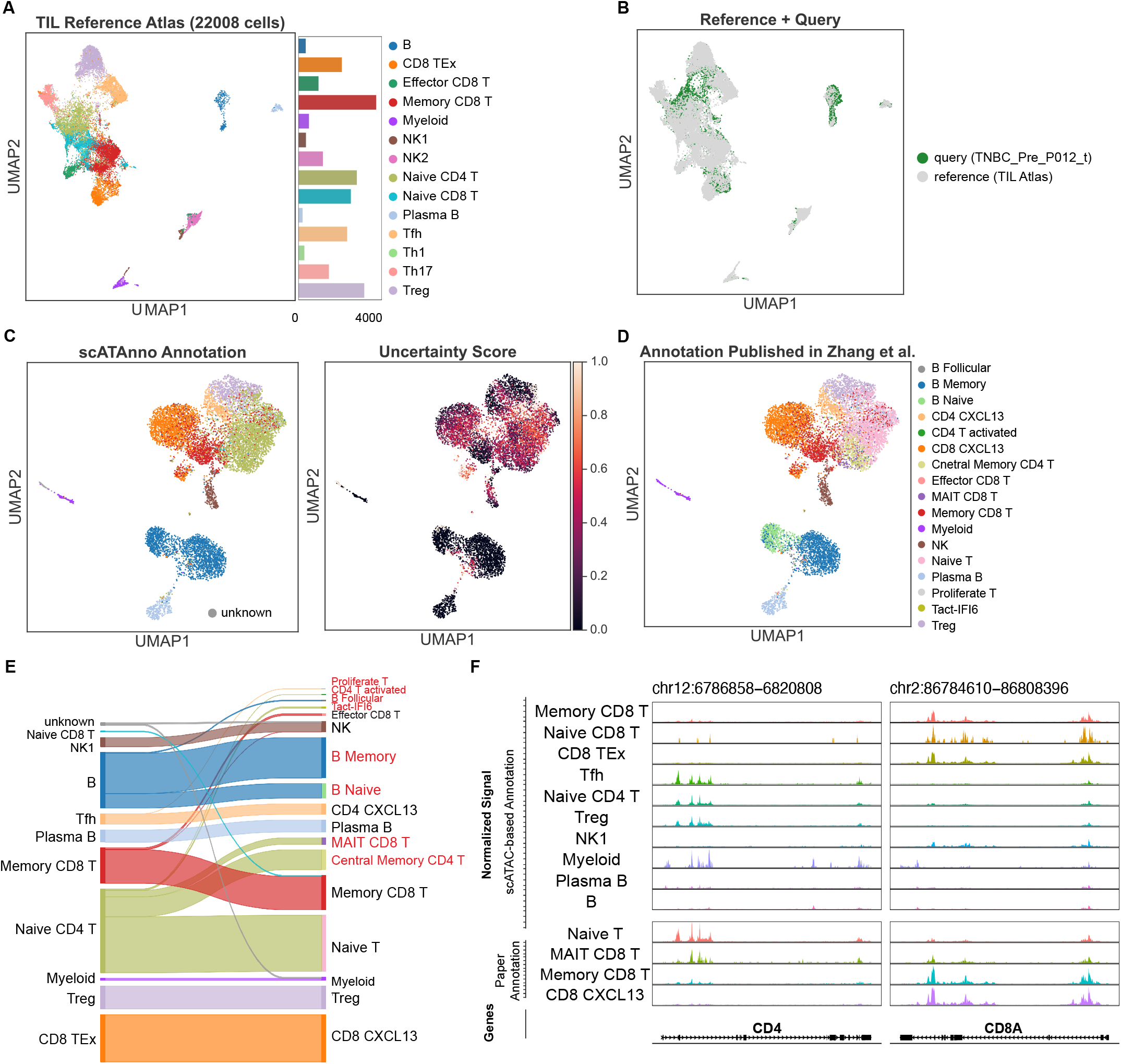
TNBC Case Study. (A) Construction of BCC TIL reference atlas (B) Integration of TNBC query data and TIL reference atlas (C) UMAP representation of query cells colored by cell types predicted from scATAnno and uncertainty score distribution (D) UMAP representation of query cells colored by cell types from the original publication (Zhang et al.) (E) Sankey plot mapping cell types annotated by scATAnno (left) to cell types annotated by the original study (right). Cell types with different names or those not represented in TIL reference atlas are highlighted in red (F) Coverage plots of chromatin accessibility over CD4 and CD8A regions across cell types predicted by scATAnno (upper panel) and across four subtypes selected from the original paper (lower panel)

### scATAnno enables annotation of tumor-infiltrating lymphocyte cell populations in the tumor microenvironment

To better characterize the subtypes of tumor-infiltrating lymphocytes, in the second step, we extract the immune cells detected above using a healthy adult reference atlas and apply a Tumor-Infiltrating Lymphocyte (TIL) reference atlas from Basal Cell Carcinoma (BCC) to produce a more fine-grained annotation of immune cells. To build the BCC TIL reference, we obtained fragment files of 17 BCC samples from raw FASTQ files [13] and generated a reference peak set (344492 peaks) by using MACS2 peaks (methods). The immune cells from two BCC query samples are well-integrated with the TIL reference data (Figure 3C-D), and most of them are accurately annotated by scATAnno (Figure 3E-H). By comparing the original annotation (Figure 3E) with scATAnno annotation (Figure 3F), only a small group of helper 1 T (Th1) cells is annotated as unknown and exhibits a high-level uncertainty score. These Th1 cells lie in the transition region in the UMAP representation, which could lead to high uncertainty scores (Figure 3F right). The remaining cells show consistent annotation results compared to the original paper annotation. Overall, scATAnno achieves an accuracy of 92.9% and a weighted F1 of 97.1%. To further confirm the TIL cell types yielded by scATAnno, we inspected the chromatin accessibility of signature genes. Each cell type exhibits enhanced levels of accessibility over signature genes (Supplementary Figure 5D). For example, the enrichment of peak signal at the PDCD1 locus, which is a T cell exhaustion signature, confirms the identity of the CD8 TEx group. And the unique accessibility in the FOXP3 locus validates the Treg cell identity. Taken together, we believe scATAnno provides robust multi-layer annotation for tumor samples. It first identifies the coarse level of cell-type annotation using the healthy adult reference and detects the immune cells in TME, and then classifies a broad spectrum of TIL subtypes of immune cells.

Overall, scATAnno not only accurately classifies cell types, but also robustly enables cross-condition cell-type annotation. We believe scATAnno can be useful for annotating scATAC-seq data under varieties of conditions.

### scATAnno leverages a tumor-infiltrating lymphocyte atlas of basal cell carcinoma to accurately predict tumor-infiltrating cells in TNBC

We demonstrate scATAnno can be applied to more complex systems, such as biopsy samples from cancer patients, and capture cellular heterogeneity of tumor-infiltrating immune cells in the tumor microenvironment (TME). We applied the TIL reference atlas from BCC samples to annotate TIL cells from a TNBC cohort [14]. To test scATAnno, we used a TNBC sample comprising 9,935 cells as query data. As shown in Figure 4A-B, scATAnno builds a TIL reference atlas encompassing 14 cell types in the UMAP space and successfully integrates the TNBC query data with the TIL reference atlas. To evaluate the performance of scATAnno, we compare the predicted query cell types (Figure 4C) with the original paper [14] which annotated the scATAC-seq cells by transferring the label from the matching scRNA-seq profiles (Figure 4D). The Sankey plot in Figure 4E compares the labels yielded by scATAnno (left) to those obtained from the original paper (right). The cell types produced by the original paper and not represented in the BCC TIL reference atlas are highlighted in red. Most cell types annotated by scATAnno are in line with the original paper. For example, B cells predicted by scATAnno are annotated as B memory, B naive, and B follicular cells in the original paper. CD8 TEx is an exhausted T cell group annotated as CD8 CXCL13 in the original study, which confirms the exhaustion features in this cell group (Yuanyuan Zhang et al. 2021). Follicular helper T (Tfh) is originally annotated as CD4 CXCL13, which has Tfh features according to the original study [14]. The cell-type annotation result can be further validated with enhanced accessibility of signature genes (Supplementary Figure 6).

We next investigate the inconsistent annotations and validate our results by examining the accessibility near genomic regions of signature genes across cell types. A group of Naive CD4 T cells is annotated as a mixture of Naive T, MAIT CD8 T, and Central Memory CD4 T in the original publication [14]. Central memory CD4 T is not represented in the BCC TIL reference atlas and hence this cell type could not be produced by scATAnno. To investigate the cell identity of originally annotated MAIT CD8 T cells, we plot the peak enrichment of MAIT CD8 T cells near the CD4 and CD8A locus and find that this group of cells exhibits strong enrichment of peaks over the CD4 than CD8A locus. Moreover, peak signals over CD4 and CD8A locus across four cell types yielded by the original study reveals a resemblance between MAIT CD8 T and Naive CD4 T cells, rather than the other CD8 T cell groups (Figure 4F). Therefore, we conclude that cells labeled as MAIT CD8 T cells are more likely CD4 T cells, as inferred by scATAnno, instead of CD8 T cells. In total, these observations demonstrate that scATAnno is a powerful tool where one can use a TIL atlas from one cancer type to identify tumor-infiltrating lymphocytes from a different cancer type, which greatly facilitates the use of scATAC to study TIL cell subtypes under complex disease conditions.

## Discussion

scATAnno is a cell-type annotation tool specifically designed for single cell chromatin accessibility data. It directly annotates new scATAC-seq data, eliminating the need for matching scRNA-seq profiling. Unlike many existing methods, scATAnno does not rely solely on the promoters and nearby regions to convert peaks into gene activity scores. Instead, it incorporates a significant number of enhancers, regardless of distance from promoter and gene body regions, providing a more comprehensive understanding of the regulatory landscape and significantly improving the characterization of cell type-specific accessibility patterns.

scATAnno enables the generation of scATAC-seq-based reference atlases from publicly available datasets and efficiently annotates cell types of newly generated query data using fragment files. Importantly, scATAnno incorporates KNN-based and weighted distance-based uncertainty scores to assess the confidence of cell type assignments and detect unknown cell populations in the query data. The novel quantification of the weighted distance uncertainty score greatly improves the accuracy of identifying unknown cells of the query data. Furthermore, scATAnno supports both single-reference atlas annotation for less complex query samples, such as PBMC, and multi-atlas annotation for samples with greater complexity.

scATAnno exhibits a strong capability of cell-type annotation at the scATAC level. In the PBMC case study, scATAnno builds a PBMC reference and improves annotation performance by correcting cells mislabeled by Seurat v3. In the TNBC study, scATAnno successfully identifies TIL subtypes in TNBC using a BCC TIL reference. In the BCC study, scATAnno develops two-round annotation tasks and shows robust cross-condition annotation using a healthy adult reference to annotate cells within the tumor microenvironment (TME). Furthermore, the scATAnno workflow can be extended to any data, allowing for reference building and query annotation beyond the presented cases.

There are limitations of scATAnno that require further development. Currently, it supports scATAC-seq data mapped to human reference genome. Future improvements aim to support different reference genome versions, such as mouse genomes. Additionally, the scope of cell types in the current reference atlases may be incomplete, resulting in the absence of some cell types using scATAnno. Moreover, while scATAnno captures the heterogeneity of most TIL subtypes in the TNBC data using a BCC TIL atlas, it has not been tested for annotating the scATAC-seq data from other cancer types. To address these limitations, we seek to incorporate more diverse atlases in the future. The rapid expansion of scATAC-seq data from multiple species and diverse systems will rapidly fill gaps in our reference sets and improve the capabilities of scATAnno and ultimately improve our understanding of chromatin configurations underling cell states in complex biological systems.

## Data and Code Availability

All datasets are publicly available. The PBMC and BCC reference data used in this study are available in the Gene Expression Omnibus (GEO) dataset under accession code GSE129785. We downloaded raw FASTQ files from BioProject (PRJNA532774), which could be found under the accession code above. The healthy adult reference data is available in the GEO dataset under accession code GSE184462. The PBMC 10K data is available in the 10X genomics datasets. We downloaded the raw data from 10X Genomics:(https://support.10xgenomics.com/single-cell-multiome-atac-gex/datasets/1.0.0/pbmc_granulocyte_sorted_10k). The TNBC data is available in the GEO dataset under accession code GSE169246. We specifically requested the original fragment file of scATAC-seq data from the first author [14]. scATAnno code is available on GitHub https://github.com/Yijia-Jiang/scATAnno-main. The ReadDocs of scATAnno tutorial and processed reference atlas and query download is available https://scatanno-main.readthedocs.io.

## Acknowledgement

The work was conducted with support from NIH grant P01 CA163227-06A1 and P01 CA250959-01, and Gray Foundation BRCA Precancer Atlas. Zhirui Hu acknowledges support from NIMH grant R01MH123178 to Katherine S. Pollard.

## Methods

### Construction of Reference Atlases

The healthy adult reference atlas is obtained from a publicly available dataset GSE184462 [12]. To build a ready-to-use atlas that first the scATAnno workflow, we obtain the count matrix provided by the original study, and selected deep-sequenced 1000 cells per minor cell type (total of 100,158 cells) and 890,130 adult-specific peaks. T cells, B cells and Myeloids are labeled as immune cells in the ready-to-use atlas.

The PBMC reference atlas is obtained from publicly available data [13]. To construct a ready-to-use atlas, we downloaded raw FASTQ files from BioProject (PRJNA532774) to generate BAM files and fragment files. The original study encompasses both bone marrow cells and PBMC cells. We excluded the bone marrow cells and selected 39,441 PBMC cells. Fragment files are concatenated and filtered by these cell barcodes using QuickATAC filter-barcodes. Peaks are called on BAMfiles using MACS2 to generate a list of 196,618 reference peaks. Next, we use QuickATAC to intersect the fragment reads on reference peaks using quick agg-countmatrix function [31]. We find that QuickATAC is unable to retain all reference peaks if there is an absence of reads within a peak location. To ensure all the reference peaks are generated in the final count matrix, we created a spike-in fake fragment file that covers the maximum and minimum chromosome start and end regions of the reference peaks, we then concatenated the spike-in fake fragment file and real fragment files to call agg-countmatrix function. QuickATAC gives output composed of MTX, barcodes, and feature CSV files. We read these files into Python-based Scanpy and construct a peak-by-cell reference matrix in Anndata format, retaining the entire peak space. We also examined chromatin accessibility over TRAV1-2, SLCA410, and PRSS35 genes, which are known as signature genes for MAIT cell type [23, 24], in the reference atlas. The original gamma delta T cell type is corrected as MAIT.

The BCC TIL atlas is constructed in a similar way as the PBMC reference. We downloaded raw FASTQ files from the same BioProject (PRJNA532774) to generate BAM files and fragment files of 17 BCC samples. We then selected 22,008 TIL cells whose cell barcodes are provided in the original study [13]. Fragment files are concatenated and filtered by these cell barcodes using QuickATAC filter-barcodes. A reference peak set (344,492 peaks) is generated by calling MACS2 on BAM files. To retain all reference peaks, we created a spike-in fragment file that covers the maximum and minimum genomic regions of the TIL peaks and then concatenated the spike-in fragment and actual fragment files to construct a ready-to-use peak-by-cell BCC TIL reference matrix.

Additionally, users can build their own reference atlases. A fragment file and a list of valid cell barcodes are required for reference building. To build a ready-to-use reference atlas, we first call peaks from the fragment file using MACS2 peak-calling algorithm [13, 16], then construct a peakby-cell count matrix of reference atlas using QuickATAC agg-countmatrix function [31]. This reference building process can be easily extended to any scATAC-seq datasets. A full script of reference building steps can be found on Github (https://github.com/Yijia-Jiang/scATAnno-main/tree/main/prep_data/scATAnno-reference-building-example).

### Query Data Preparation

PBMC cells isolated through Fluorescence-activated cell sorting (FACS) data is downloaded from GSE123578. We downloaded raw fragment files mapped to hg19 reference genome from GEO database, and lift over the fragments mapping to hg38 genome. The lymph node tumor of a patient diagnosed with B cell Lymphoma can be downloaded from 10X(https://www.10xgenomics.com/resources/datasets/). And the ground truth label is provided by 10X (https://pages.10xgenomics.com/rs/446-PBO-704/images/10x_LIT000110_Data_Spotlight_Multiome_digital.pdf).

The original dataset has low quality cells and unambiguous cell type labels, such as mixed T and B cells. To prepare a clean dataset for benchmarking, we removed cells with low quality and curated the cell type labels for B cells, Tumor B cells, T cells and Monocytes.

PBMC 10K is directly downloaded from 10X Genomics:(https://support.10xgenomics.com/single-cell-multiome-atac-gex/datasets/). To align the PBMC 10K features with the PBMC reference peaks, we concatenated the fragment file of PBMC 10K with PBMC spike-in fragment file, and use QuickATAC agg-countmatrix function to produce the MTX, barcodes and feature CSV files. A final peak-by-cell matrix of query data is constructed using Scanpy [22]. The Seurat v3 annotation is directly obtained from calling “LoadData(“pbmcMultiome”, “pbmc.rna”)” function in R, as shown in Vignette (https://satijalab.org/seurat/articles/atacseq_integration_vignette.html). We curated some cell type names from Seurat annotation to make the cell labels more comparable. Modifications include merging CD14 and CD16 monocytes as Monocytes, combining Effector and Central memory CD4 T as Memory CD4 T cells, and merging CD8 effector memory T1 and T2 as Effector memory CD8 T cells.

The TNBC fragment file is requested from the first author of the TNBC paper [14]. To align the TNBC features with the TIL reference peaks, we concatenated the fragment file of TNBC with TIL spike-in and used QuickATAC agg-countmatrix and Scanpy to generate the peak-by-cell matrix of TNBC data.

BCC fragment files are held out from the BCC TIL reference building [13]. To align the BCC peaks with the TIL reference peaks, we concatenated the fragment file of BCC with TIL spike-in fragment reads, and used QuickATAC agg-countmatrix and Scanpy to generate the peak-by-cell matrix of BCC data. Optionally, QuickATAC provides quality control metrics, such as TSS enrichment scores, and users can set criteria to filter out low-quality cells.

### Dimensionality Reduction

To reduce the high dimensionality of scATAC-seq data, we incorporated SnapATAC2 to perform spectral embedding [12, 17]. The reference and query data are concatenated to form a combined peak-by-cell matrix. SnapATAC2 transforms the count matrix to a cell-cell similarity matrix using the Jaccard Similarity Score, which computes the ratio of the intersection and the union of two cells to estimate the similarity between cells [17]. It then normalizes the Jaccard similarity matrix to regress out the sequencing depth bias. Next, it computes the normalized Jaccard similarity matrix followed by eigenvector decomposition to obtain the low dimensional components. To reduce the computational cost of analyzing high dimensional data, SnapATAC2 first sampled “landmark” cells that capture the data distribution of the original points, and obtain the low dimensional components of these highly informative cells. It finally projects the remaining cells into the same low-dimensional components and obtains low-dimensional components for all cells [2, 17].

### Batch Correction

To account for the potential technical variability of the reference atlas and query data, we apply the batch correction method, Harmony, to eliminate any potential batch effects [20]. Harmony is applied to low-dimensional embeddings of the concatenated reference and query dataset to regress out batch effects. The harmonized embeddings better capture the true biological signals of the reference atlas and query data.

### Cell-type assignment

In the initial cell-type assignment, each query cell is assigned a cell label using its closest K-nearest neighbors (KNN) in the reference atlas, based on the low dimensional embeddings. The KNN algorithm is implemented using the Scikit-learn package in Python [32].

### Uncertainty Score Measurement

Suppose 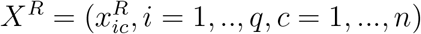, a *q × n* matrix, is the coordinates of *n* reference cells in the latent space, each column is a cell and each row is a coordinate.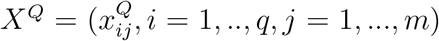, a *q × m* matrix, is the coordinates of *m* query cells.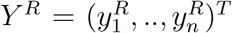, a *n*-dim vector, each 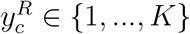 is the cell type label of each reference cell; 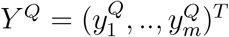, a *m*-dim vector, each 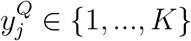 is the predicted cell type label.

#### KNN-based Uncertainty Score

In the first step, we compute the uncertainty score of cell type assignment of each query cell. For query cell *j*, identify *p* nearest reference cells (*𝒩*_*j*_) based on Euclidean distance of low dimension embeddings, i.e. dist 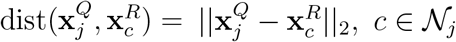 . We computed the standard deviation of these nearest distances: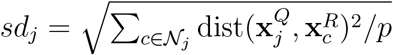 . Then, we apply Gaussian kernel function to transform the distance to a similarity measurement: 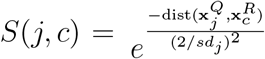, *c ∈ 𝒩*_*j*_ . Finally, we compute the probability of assigning cell type to each query cell: 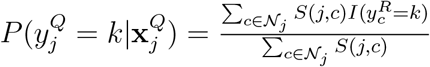 . Then the uncertainty score is 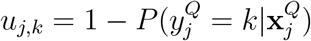 . Based on it, we can assign cell types:

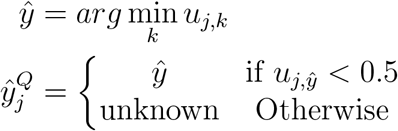

and the uncertainty score for cell *j* in the first step is:

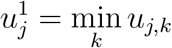

#### Weighted Distance-Based Uncertainty Score

The goal of computing the Weighted Distance-Based uncertainty score is to measure the distance of each query cell and the cell type assigned to it, with an intuition that a query cell far away from the assigned cell type cluster is likely to be an unknown cell. The computation of the Weighted Distance-Based Uncertainty Score takes two steps: 1) compute the weights of low dimensional embeddings for a specific cell type k to capture the relative importance in discriminating cell type k, and 2) compare within cell-type reference distance and the weighted distance of each query cell and the centroid of its assigned cell type k.

In the first step, we compute the weight of each latent dimension to distinguish each cell type from the rest, based on the assumption that low dimensional components have different discriminate power for a specific cell type. For cell type k, the weight of each dimension *i, W*_*ik*_, *i* = 1, …, *q*, is defined as the ratio of average between cell-type distances and within cell-type variation, which is computed using n reference cells as follows:

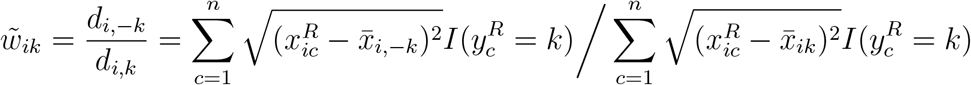

where 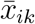 is the robust centroid of cell type *k* after removing outlier cells, and 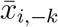 is the robust centroid of the rest of cells except cell type *k*.

Specifically, we first identify the centroid of cell type k and the centroid of the remaining cells at ith embedding. We first remove the outlier cells based on UMAP to construct a cleaner structure of reference atlas and then compute weights. We compute within cell-type distance d(i,k), which is the distance of reference cells of cell type k and reference centroid of cell type k, and compute between cell-type distance d(i,-k), which is the distance of reference cells of cell type k and the centroid of remaining cell types (Formula 1). The ratio of average between cell-type distances and within cell-type distances is the weight of embedding i for cell type k. We repeat the process to get the weights of all latent embeddings for cell type k and normalize the weights. *W*_*ik*_ is higher if reference cells are closer to their respective cell type centroid and far away from the other cell types. And embeddings assigned with high weights are more important to distinguish a given cell type.

Then, we normalized 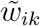 for each cell type:

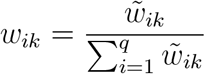

In the second step, based on these weights, we compute within cell-type weighted distance for reference cells and compare the distance of each query cell to these reference distances. If the distance of the query cell is significantly larger than the distance of reference cells, we reject the annotation of the query cell. For query cell *j* with predicted cell type label 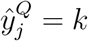, we compute the weighted distance for each reference cell in cell type *k* and for the query cell:

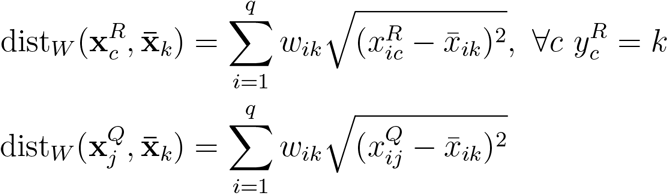

Then, we compare the weighted distance of the query cell to 95% quantiles of all reference cells with cell type *k*. If the query distance is greater than 95 percentile of the within-cluster distance of cell type *k*, we set the uncertainty score to 1.0 and annotate the cell as unknown:

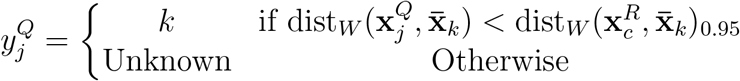

and the uncertainty score for cell *j* in the second step is:

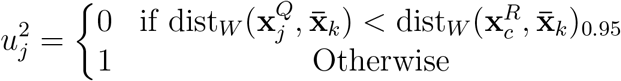

### Cluster-level Annotation

If no prior cluster information is provided, we cluster query cells using the Leiden algorithm to assign similar cells into the same group based on low-dimensional embeddings. The Leiden algorithm is implemented using scanpy.tl.leiden and applied to the query data with high resolution [22, 33]. We determine the cell type of each cluster by taking the most abundant cell type. If cluster information is provided by users, scATAnno uses the cluster information and assigns the cluster-level annotation by majority voting, i.e., taking the most abundant cell type. In this way, all query cells are annotated at the cluster-level annotation.

### Coverage Plots of scATAC-seq Data

Coverage plots are generated using Signac [24]. First, the fragment index file is generated from the fragment file for each query data. Then, the fragment file is read into Signac and the accessibility of peaks over genomic regions is plotted using the CoveragePlot function in Signac [24].

## Figure Captions

## Supplementary Figures

Supplementary Figure 1: Cell type annotation of holdout healthy adult cells. (A) Construction of Healthy Adult reference atlas (B) Integration of holdout healthy adult cells and the reference atlas (C) UMAP representation of query cells annotated by original publication (Sapathy et al.) (D) UMAP representation of query cells colored by cell types and uncertainty score distribution produced by scATAnno (E) Computation of F1 score (left) and accuracy (right) of each cell type (F) Bar plot of average accuracy scores of using promoter regions (extending 1kb, 2kb, 5kb, 10kb) and all peak regions. Overall accuracy scores are 3.37%, 34.2%, 58.3%, 77.2% and 98.2% respectively. (G) Box plot of accuracy scores breaking down by each cell type using promoter regions (extending 1kb, 2kb, 5kb, 10kb) and all peak regions. Each color dot shows the accuracy score of a cell type shown in the legend.

Supplementary Figure 2: Simulation of annotating holdout healthy adult cells using ablated Reference Atlas. (A) Healthy Adult reference atlas using all reference cells highlighting Vascular Smooth cells and Adrenal Cortical cells (B) Construction of a new Healthy Adult reference atlas ablated Vascular Smooth cells from the original atlas (C) Construction of a new Healthy Adult reference atlas ablated Adrenal Cortical cell type (D) Query cells on UMAP representation, highlighting vascular smooth cells in a dashed circle. The arrow points to the three-step annotation results for the Vascular Smooth cells: KNN assignment, weighted distance-based assignment, and clusterlevel assignment (E) Proportion of correctly identifying Vascular Smooth as unknown cells in each assignment step. Proportion is computed as the number of unknown cells divided by the total number of vascular smooth cells in the query data. (F) Query cells on UMAP representation, highlighting Adrenal Cortical cells in a dashed circle and three-step annotation results (G) Proportion of correctly identifying Adrenal Cortical as unknown cells in each assignment step, following the same computation as (E)

Supplementary Figure 3: PBMC Case Study. (A) Constructed PBMC reference atlas (B) Integration of PBMC query data and the reference atlas (C) Coverage plots of chromatin accessibility over TRAV1-2, SLC4A10, and PRSS35 genomic regions across cell types predicted by scATAnno

Supplementary Figure 4: Benchmarking. A-D) benchmark B lymphoma data. A) UMAP of ground truth cell labels of B cell lymphoma data, compared with cell type labels yielded by scATAnno, Cellcano and Signac. B) Accuracy scores of seven methods (scATAnno, Cellcano, scJoint, Signac, Seurat, singleR and scID) for B cell lymphoma annotation. C) F1 scores of seven methods for B cell lymphoma annotation. D) Proportion of tumor B cells correctly identified as unknown. E) Bar plot of accuracy scores of scATAnno, Cellcano, scJoint, Signac, Seurat, singleR and scID for FACS-sorted PBMC data in five cell type annotation tasks (4 ablation tasks and one task using all reference cells). F) Box plot of proportion of cells correctly identified as unknown in PBMC in 4 ablation tasks. G) Heatmap of accuracy scores for five annotation tasks across methods. Four ablation scenario of removing one celltype from the reference atlas: remove B cells, remove CD4 T cells, remove CD8 T cells, remove Monocytes and remove NK cells. One task uses all reference cells.

Supplementary Figure 5: (A) Construction of Healthy Adult reference atlas (B) Integration of two holdout BCC samples and healthy reference atlas (C) Uncertainty score of query cells visualized in the integrated UMAP (D) Coverage Plots of BCC Case Study. Each plot shows the chromatin accessibility of a signature gene across cell types. The cell types are obtained from the combination of two-round annotations. Chromatin accessibility of the CD3D locus is enriched for T cells, and CD8A and CD4 loci are enriched for CD8 T and CD4 cells respectively. Chromatin accessibility of PDCD1 enriched for CD8 TEx and Tfh, and chromatin accessibility of FOXP3 enriched for Treg. Chromatin accessibility of GNLY enriched for NK, S100A9 enriched for myeloid, PECAM1 enriched for Endothelial cells. Chromatin accessibility of CD79A is enriched for B cells, and CD27 is enriched for Plasma B cells. Chromatin accessibility of PDGFRB is enriched for Mural cells.

Supplementary Figure 6: Coverage Plots of TNBC Case Study. Each plot shows the chromatin accessibility of a signature gene across predicted cell types. Chromatin accessibility of CD3D locus is enriched for T cells, chromatin accessibility of FOXP3 enriched for Treg, chromatin accessibility of PDCD1 enriched for CD8 TEx and Tfh, chromatin accessibility of GNLY enriched for NK, chromatin accessibility of CCR7 enriched for Naive T cells, chromatin accessibility CXCR5 enriched for Tfh, chromatin accessibility of S100A9 enriched for myeloid, chromatin accessibility of CD79A enriched for B cells and CD27 enriched for Plasma B cells.

## References

[1] J. D. Buenrostro, B. Wu, U. M. Litzenburger, D. Ruff, M. L. Gonzales, M. P. Snyder, H. Y. Chang, W. J. Greenleaf, Single-cell chromatin accessibility reveals principles of regulatory variation, Nature 523 (7561) (2015) 486–490.

[2] S. Preissl, K. J. Gaulton, B. Ren, Characterizing cis-regulatory elements using single-cell epigenomics, Nat. Rev. Genet. 24 (1) (2023) 21–43.

[3] D. Aran, A. P. Looney, L. Liu, E. Wu, V. Fong, A. Hsu, S. Chak, R. P. Naikawadi, P. J. Wolters, A. R. Abate, A. J. Butte, M. Bhattacharya, Reference-based analysis of lung single-cell sequencing reveals a transitional profibrotic macrophage, Nat. Immunol. 20 (2) (2019) 163–172.

[4] C. Xu, R. Lopez, E. Mehlman, J. Regier, M. I. Jordan, N. Yosef, Probabilistic harmonization and annotation of single-cell transcriptomics data with deep generative models, Mol. Syst. Biol. 17 (1) (2021) e9620.

[5] H. A. Pliner, J. Shendure, C. Trapnell, Supervised classification enables rapid annotation of cell atlases, Nat. Methods 16 (10) (2019) 983–986.

[6] A. Piovesan, F. Antonaros, L. Vitale, P. Strippoli, M. C. Pelleri, M. Caracausi, Human protein-coding genes and gene feature statistics in 2019, BMC Res. Notes 12 (1) (2019) 315.

[7] ENCODE Project Consortium, J. E. Moore, M. J. Purcaro, H. E. Pratt, C. B. Epstein, N. Shoresh, J. Adrian, T. Kawli, C. A. Davis, A. Dobin, R. Kaul, J. Halow, E. L. Van Nostrand, P. Freese, D. U. Gorkin, Y. Shen, Y. He, M. Mackiewicz, F. Pauli-Behn, B. A. Williams, A. Mortazavi, C. A. Keller, X.-O. Zhang, S. I. Elhajjajy, J. Huey, D. E. Dickel, V. Snetkova, X. Wei, X. Wang, J. C. Rivera-Mulia, J. Rozowsky, J. Zhang, S. B. Chhetri, J. Zhang, A. Victorsen, K. P. White, A. Visel, G. W. Yeo, C. B. Burge, E. Lécuyer, D. M. Gilbert, J. Dekker, J. Rinn, E. M. Mendenhall, J. R. Ecker, M. Kellis, R. J. Klein, W. S. Noble, A. Kundaje, R. Guigó, P. J. Farnham, J. M. Cherry, R. M. Myers, B. Ren, B. R. Graveley, M. B. Gerstein, L. A. Pennacchio, M. P. Snyder, B. E. Bernstein, B. Wold, R. C. Hardison, T. R. Gingeras, J. A. Stamatoyannopoulos, Z. Weng, Expanded encyclopaedias of DNA elements in the human and mouse genomes, Nature 583 (7818) (2020) 699–710.

[8] T. Stuart, A. Butler, P. Hoffman, C. Hafemeister, E. Papalexi, W. M. Mauck, 3rd, Y. Hao, M. Stoeckius, P. Smibert, R. Satija, Comprehensive integration of Single-Cell data, Cell 177 (7) (2019) 1888–1902.e21.

[9] W. Ma, J. Lu, H. Wu, Cellcano: supervised cell type identification for single cell ATAC-seq data, Nat. Commun. 14 (1) (2023) 1864.

[10] C. Wang, D. Sun, X. Huang, C. Wan, Z. Li, Y. Han, Q. Qin, J. Fan, X. Qiu, Y. Xie, C. A. Meyer, M. Brown, M. Tang, H. Long, T. Liu, X. S. Liu, Integrative analyses of single-cell transcriptome and regulome using MAESTRO, Genome Biol. 21 (1) (2020) 198.

[11] H. Chen, C. Lareau, T. Andreani, M. E. Vinyard, S. P. Garcia, K. Clement, M. A. Andrade-Navarro, J. D. Buenrostro, L. Pinello, Assessment of computational methods for the analysis of single-cell ATAC-seq data, Genome Biol. 20 (1) (2019) 241.

[12] K. Zhang, J. D. Hocker, M. Miller, X. Hou, J. Chiou, O. B. Poirion, Y. Qiu, Y. E. Li, K. J. Gaulton, A. Wang, S. Preissl, B. Ren, A single-cell atlas of chromatin accessibility in the human genome, Cell 184 (24) (2021) 5985–6001.e19.

[13] A. T. Satpathy, J. M. Granja, K. E. Yost, Y. Qi, F. Meschi, G. P. McDermott, B. N. Olsen, M. R. Mumbach, S. E. Pierce, M. R. Corces, P. Shah, J. C. Bell, D. Jhutty, C. M. Nemec, J. Wang, L. Wang, Y. Yin, P. G. Giresi, A. L. S. Chang, G. X. Y. Zheng, W. J. Greenleaf, H. Y. Chang, Massively parallel single-cell chromatin landscapes of human immune cell development and intratumoral T cell exhaustion, Nat. Biotechnol. 37 (8) (2019) 925–936.

[14] Y. Zhang, H. Chen, H. Mo, X. Hu, R. Gao, Y. Zhao, B. Liu, L. Niu, X. Sun, X. Yu, Y. Wang, Q. Chang, T. Gong, X. Guan, T. Hu, T. Qian, B. Xu, F. Ma, Z. Zhang, Z. Liu, Single-cell analyses reveal key immune cell subsets associated with response to PD-L1 blockade in triple-negative breast cancer, Cancer Cell 39 (12) (2021) 1578–1593.e8.

[15] D. A. Cusanovich, R. Daza, A. Adey, H. A. Pliner, L. Christiansen, K. L. Gunderson, F. J. Steemers, C. Trapnell, J. Shendure, Multiplex single cell profiling of chromatin accessibility by combinatorial cellular indexing, Science 348 (6237) (2015) 910–914.

[16] Y. Zhang, T. Liu, C. A. Meyer, J. Eeckhoute, D. S. Johnson, B. E. Bernstein, C. Nusbaum, R. M. Myers, M. Brown, W. Li, X. S. Liu, Model-based analysis of ChIP-Seq (MACS), Genome Biol. 9 (9) (2008) R137.

[17] R. Fang, S. Preissl, Y. Li, X. Hou, J. Lucero, X. Wang, A. Motamedi, A. K. Shiau, X. Zhou, F. Xie, E. A. Mukamel, K. Zhang, Y. Zhang, M. M. Behrens, J. R. Ecker, B. Ren, Comprehensive analysis of single cell ATAC-seq data with SnapATAC, Nat. Commun. 12 (1) (2021) 1337.

[18] GitHub - kaizhang/SnapATAC2: Single-cell epigenomics analysis tools, https://github.com/kaizhang/SnapATAC2, accessed: 2023-5-15.

[19] K. Zhang, N. R. Zemke, E. J. Armand, B. Ren, A fast, scalable and versatile tool for analysis of single-cell omics data, Nature Methods (2024) 1–11.

[20] I. Korsunsky, N. Millard, J. Fan, K. Slowikowski, F. Zhang, K. Wei, Y. Baglaenko, M. Brenner, P.-R. Loh, S. Raychaudhuri, Fast, sensitive and accurate integration of single-cell data with harmony, Nat. Methods 16 (12) (2019) 1289–1296.

[21] M. Lotfollahi, M. Naghipourfar, M. D. Luecken, M. Khajavi, M. Büttner, M. Wagenstetter, Ž. Avsec, A. Gayoso, N. Yosef, M. Interlandi, S. Rybakov, A. V. Misharin, F. J. Theis, Mapping single-cell data to reference atlases by transfer learning, Nat. Biotechnol. 40 (1) (2022) 121–130.

[22] F. A. Wolf, P. Angerer, F. J. Theis, SCANPY: large-scale single-cell gene expression data analysis, Genome Biol. 19 (1) (2018) 15.

[23] C. Domínguez Conde, C. Xu, L. B. Jarvis, D. B. Rainbow, S. B. Wells, T. Gomes, S. K. Howlett, O. Suchanek, K. Polanski, H. W. King, L. Mamanova, N. Huang, P. A. Szabo, L. Richardson, L. Bolt, E. S. Fasouli, K. T. Mahbubani, M. Prete, L. Tuck, N. Richoz, Z. K. Tuong, L. Campos, H. S. Mousa, E. J. Needham, S. Pritchard, T. Li, R. Elmentaite, J. Park, E. Rahmani, D. Chen, D. K. Menon, O. A. Bayraktar, L. K. James, K. B. Meyer, N. Yosef, M. R. Clatworthy, P. A. Sims, D. L. Farber, K. Saeb-Parsy, J. L. Jones, S. A. Teichmann, Cross-tissue immune cell analysis reveals tissue-specific features in humans, Science 376 (6594) (2022) eabl5197.

[24] T. Stuart, A. Srivastava, S. Madad, C. A. Lareau, R. Satija, Single-cell chromatin state analysis with signac, Nature methods 18 (11) (2021) 1333–1341.

[25] W. Ma, J. Lu, H. Wu, Cellcano: supervised cell type identification for single cell atac-seq data, Nature Communications 14 (1) (2023) 1864.

[26] Y. Lin, T.-Y. Wu, S. Wan, J. Y. Yang, W. H. Wong, Y. R. Wang, scjoint integrates atlas-scale single-cell rna-seq and atac-seq data with transfer learning, Nature biotechnology 40 (5) (2022) 703–710.

[27] K. Boufea, S. Seth, N. Batada, scid uses discriminant analysis to identify transcriptionally equivalent cell types across single-cell rna-seq data with batch effect. iscience 23 (3): 100914 (2020).

[28] R. Sun, X. Kong, X. Qiu, C. Huang, P.-P. Wong, The emerging roles of pericytes in modulating tumor microenvironment, Front Cell Dev Biol 9 (2021) 676342.

[29] F. Feng, X. Feng, D. Zhang, Q. Li, L. Yao, Matrix stiffness induces Pericyte-Fibroblast transition through YAP activation, Front. Pharmacol. 12 (2021) 698275.

[30] O. Franzén, L.-M. Gan, J. L. M. Björkegren, PanglaoDB: a web server for exploration of mouse and human single-cell RNA sequencing data, Database 2019 (Jan. 2019).

[31] GitHub - AllenWLynch/QuickATAC, https://github.com/AllenWLynch/QuickATAC, accessed: 2023-4-18.

[32] R. Garreta, G. Moncecchi, Learning scikit-learn: Machine Learning in Python, Packt Publishing Ltd, 2013.

[33] V. A. Traag, L. Waltman, N. J. van Eck, From louvain to leiden: guaranteeing well-connected communities, Sci. Rep. 9 (1) (2019) 5233.

