## Supplementary figures and images for "scATAnno: Automated Cell Type Annotation for single-cell ATAC Sequencing Data"

### SupplementaryFigure1_simulation.pdf

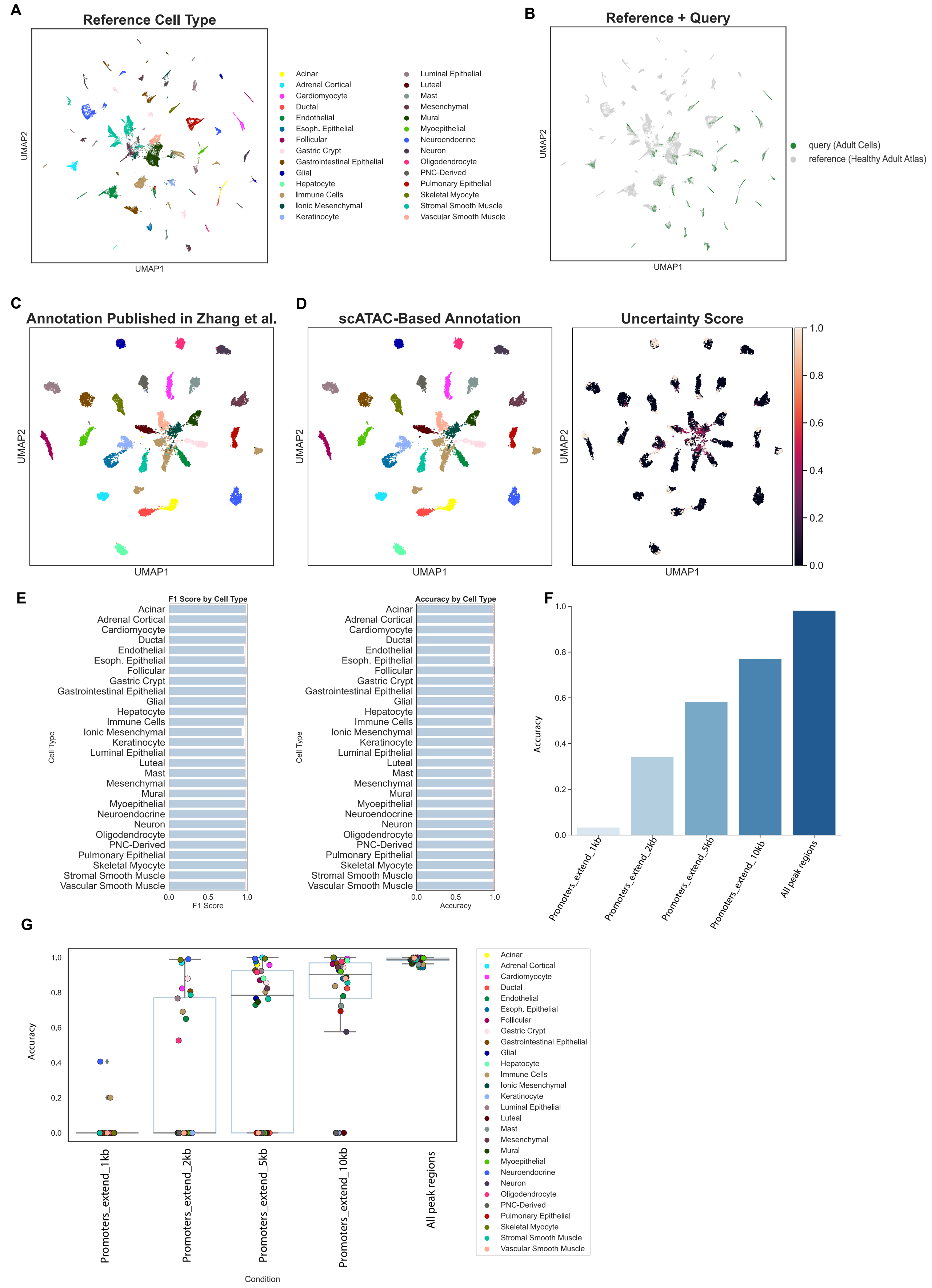


Supplementary Figure 1

### SupplementaryFigure2_ablation.pdf

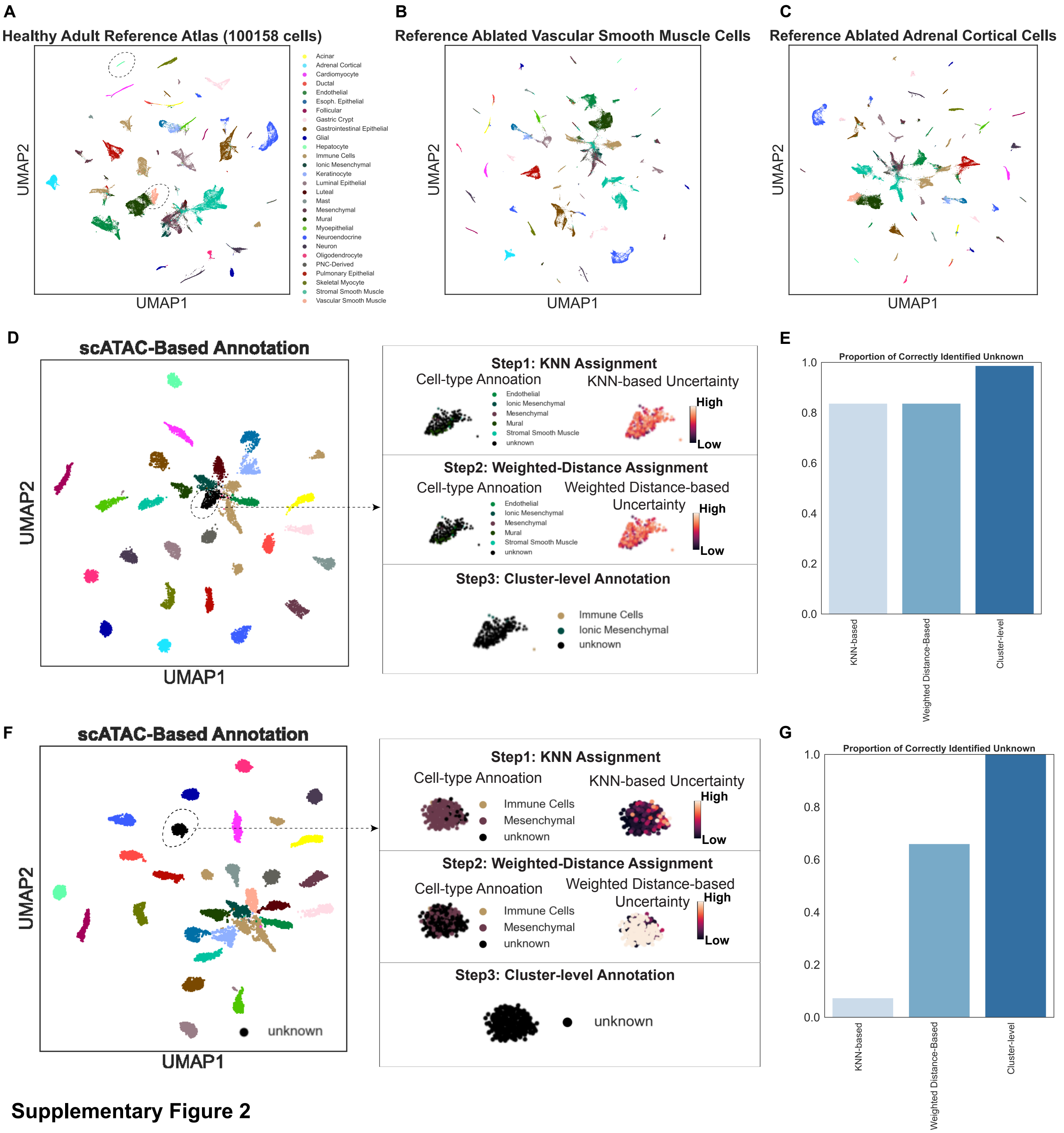


**Supplementary Figure 2**

### SupplementaryFigure3_PBMC.pdf

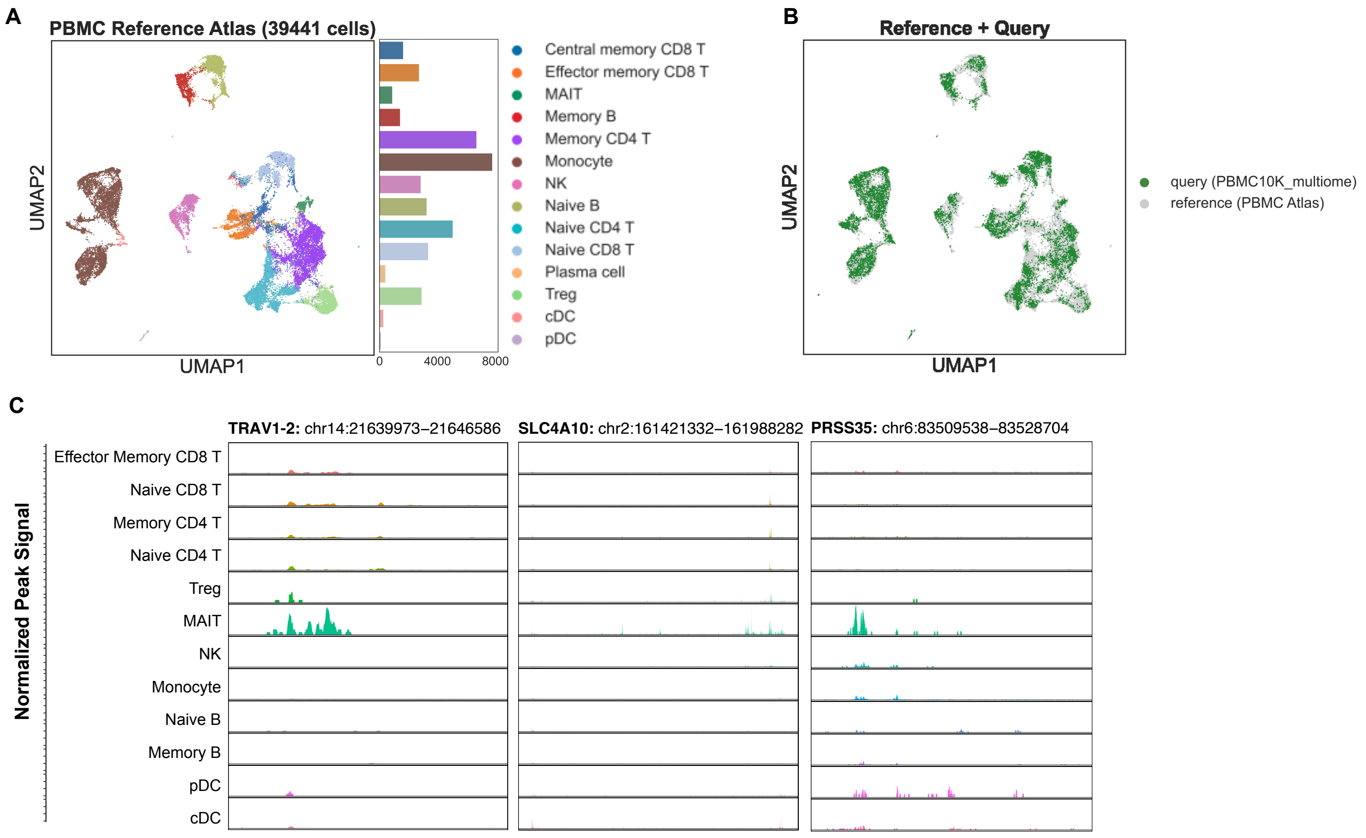


#### Supplementary Figure 3

### SupplementaryFigure4_benchmark.pdf

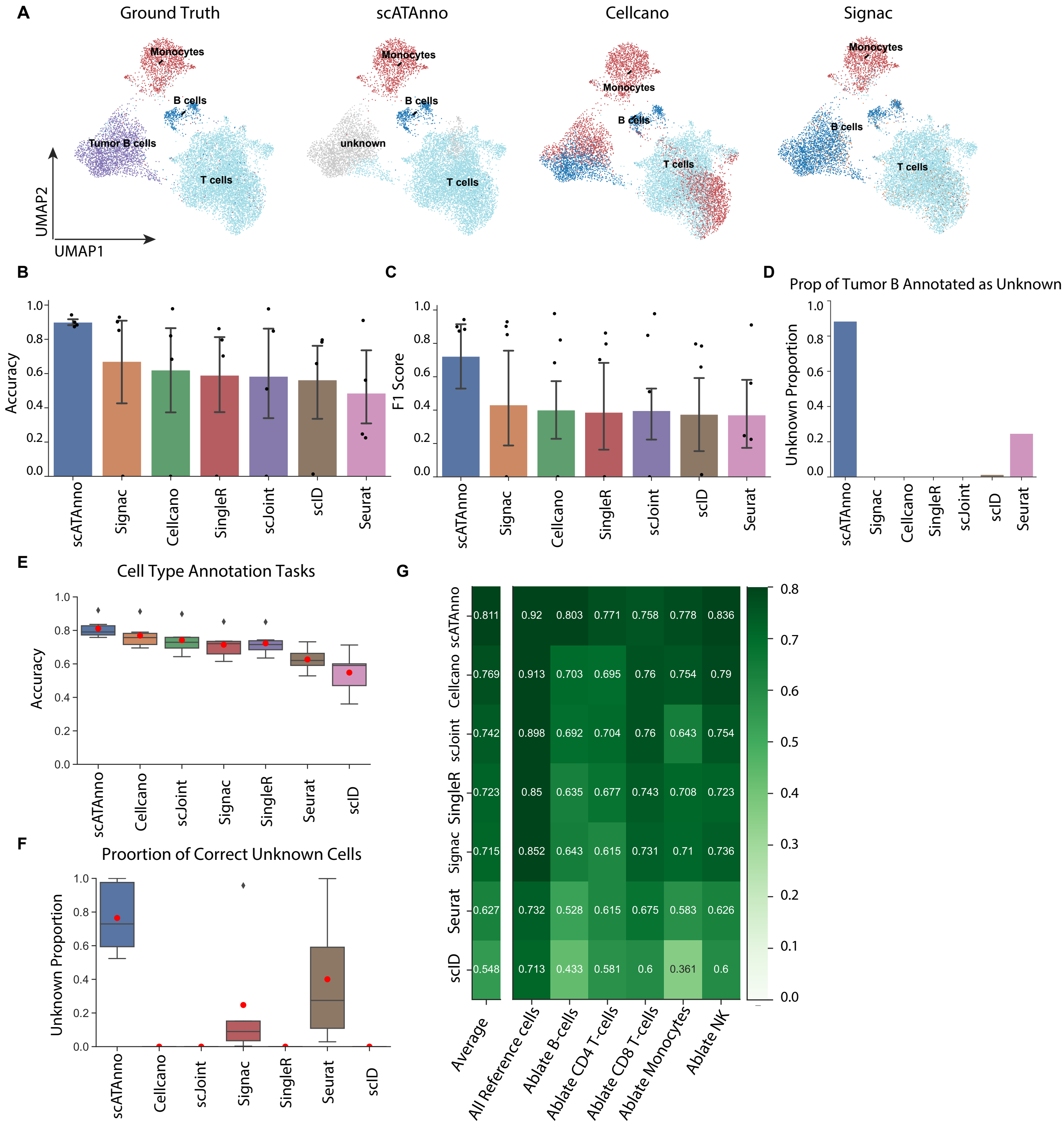


**Supplementary Figure 4**

### SupplementaryFigure5_BCC_coverage.pdf

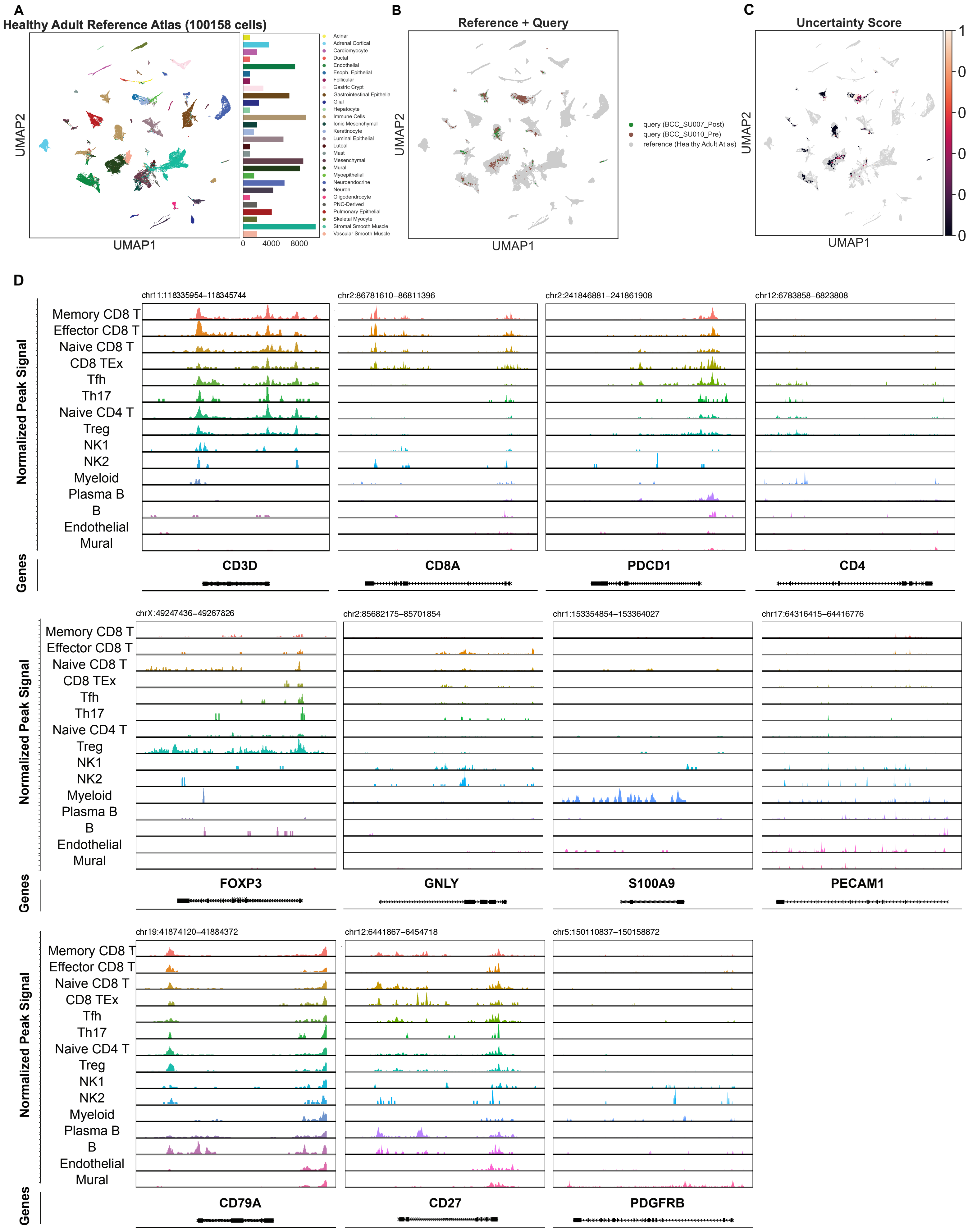


**Supplementary Figure 5**

### SupplementaryFigure6_TNBC_coveragePlot.pdf

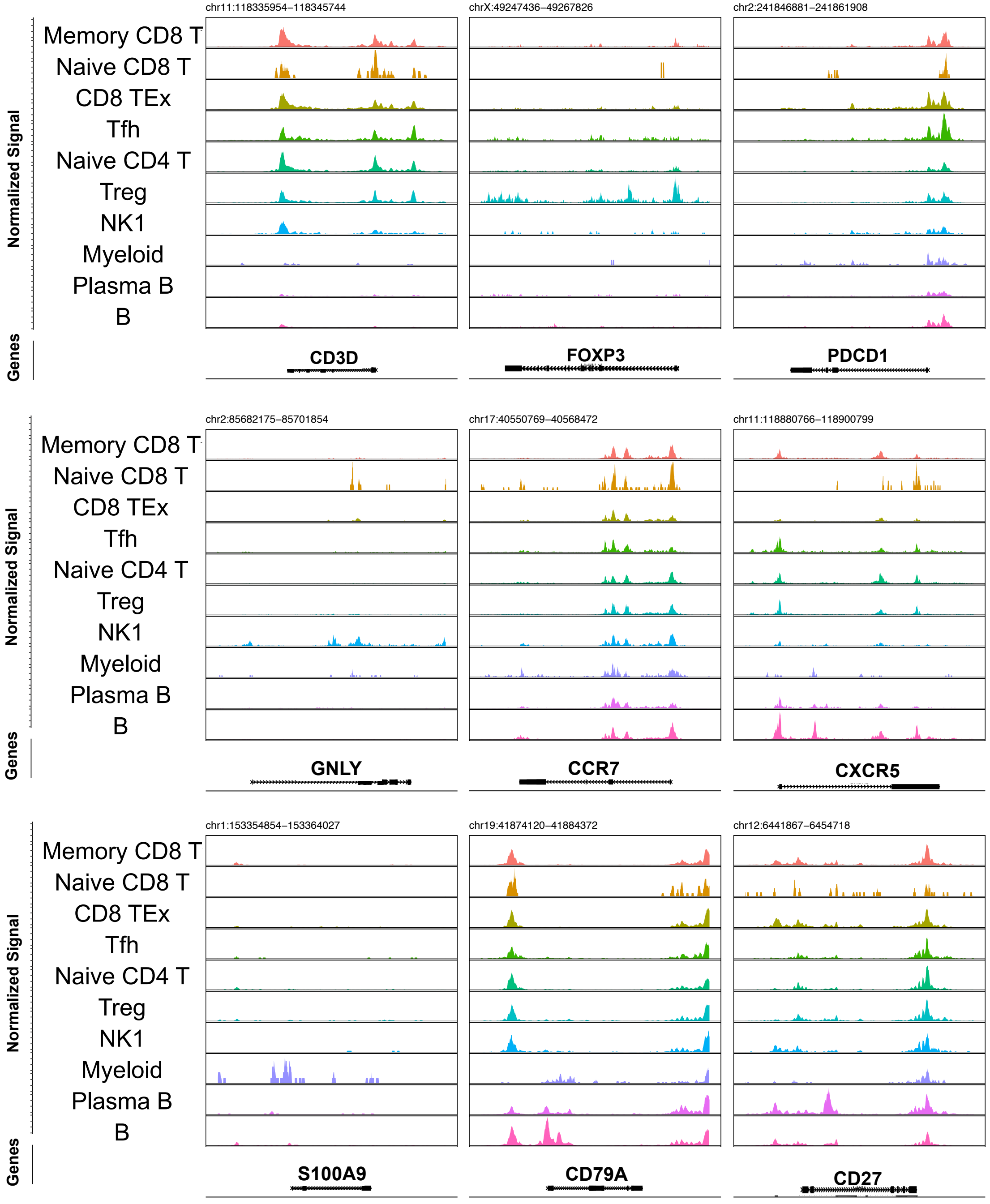


**Supplementary Figure 6**
