## Supplementary material for "scATAnno: Automated Cell Type Annotation for single-cell ATAC Sequencing Data": table 1

Table 1: Key features of scATAnno compared to other cell-type annotation methods

| Method | Version | Build for scATAC-seq Annotation | Use Distal Enhancer Signal | Do Not Need Matched scRNA-seq | Provide Uncertainty Score | Predict Unknown Cells |
|---|---|---|---|---|---|---|
| scATAnno | 1.0.0 | ✔ | ✔ | ✔ | ✔ | ✔ |
| Cellcano | 1.0.2 | ✔ | | ✔ | | |
| scjoint | 1.0.0 | | | | | |
| Seurat | 4.3.0 | | | | ✔ | ✔ |
| Signac | 1.9.0 | ✔ | | | ✔ | ✔ |
| SingleR | 0.2.2 | | | | | |
| scID | 2.0.0 | | | | | |
